# Parental allele-specific genome architecture and transcription during the cell cycle

**DOI:** 10.1101/201715

**Authors:** Haiming Chen, Sijia Liu, Laura Seaman, Cyrus Najarian, Weisheng Wu, Mats Ljungman, Gerald Higgins, Alfred Hero, Max Wicha, Indika Rajapakse

**Author notes:** These authors contributed equally (Haiming Chen, Sijia Liu). Correspondence to Indika Rajapakse and Max Wicha.

## Abstract

A normal human somatic cell inherits two haploid genomes. Individual chromosomes of each pair have distinct parental origins and parental alleles are known to unequally contribute to cellular function. We integrated chromosome conformation (form) and gene transcription (function) analyses to dissect the dynamics of the maternal and paternal genomes in lymphoblastoid cells during the cell cycle. We found a distinct set of homologous alleles with very different activity often located close to boundaries of euchromatin and heterochromatin domains. We also identified a set of allele-biased topologically associating domains (TADs) that were small sized and had higher gene density. Thousands of genes show allelically biased expression (ABE) with false discovery rate < 0.05, and 98% of them have no allelic switching during G1, S, and G2/M phases. A subset of ABE genes are preferentially localized near TAD boundaries, enriched with chromatin organization transcription factor binding sites, and contained higher number of sequence variants in CCCTC-binding factor sites. Our results extend previous findings of sequence variation as a basis for unequal functional parental genomes. Investigation of haplotype-resolved form-function dynamics may further our understanding of phenotypic traits, genetic diseases, vulnerability to complex disorders, and the development of cancers.

## Introduction

A somatic cell contains two haploid genomes, one with paternal origin (Pat), and the other with maternal origin (Mat). The two genomes are unequal in DNA sequence ^1,2^, gene expression^3,4^, chromatin state^5–7^, and architecture^8 9^. In linear DNA, millions of variants exist between the Pat and Mat genomes, including single nucleotide variants (SNVs) and short sequence inversions/deletions (indels) that make up 99.9% all types of variants ^1,10^. In terms of gene expression, the selection of an allele to be expressed leads to mono-allelic expression (MAE) or allelically biased expression (ABE). Known cases of MAE include X-linked genes due to X chromosome inactivation in females ^11^, and imprinted genes inherited from genomic imprinting ^12–15^. Recent genome wide transcriptomics analyses reveal that approximately 20% of human genes show ABE ^16–18^. The selection of an allele to be expressed appears to be stochastic and independent of parental origin ^4,19^, and monoallelic transcripts could occupy as much as 74% of the total transcripts in a cell ^19^.

With regard to chromatin states, the two genomes are unequally modified by cytosine methylation at the canonical CpG or non-CpG sites and core histone remodeling (e.g., acetylation, methylation, or ubiquitination) ^5–7,20^. In addition, homologous chromosomes are differentially organized in three dimensional (3D) structures ^8,9^. For examples, the inactive X chromosome in females is partitioned into two “super-domains” ^8^; loop structure differences are also present at imprinted loci ^8^; and during stem cell differentiation, a small number of alleles switched between euchromatin and heterochromatin states (without enrichment of ABE or known imprinted genes)^91^.

The captured variations in sequence, allele selection for expression, chromatin state and organization provide different points of view of the Pat and Mat genomes within diploid cells. However, it remains a challenge to integrate such “independent” information into a systematic view of the dynamics of individual genomes, including both chromatin organization (form) and gene transcription (function). In addition, it is unclear how ABE is maintained during the cell cycle in proliferating cells.

In this study, we take advantage of a haplotype-resolved diploid genome of the cell line NA12878 ^10^ to address these issues. We capture nascent gene transcripts with bromo-uridine labeling and sequencing (Bru-seq) ^21^, cytoplasmic mRNA species with RNA sequencing (RNA-seq)^22^, and chromatin interactions with Hi-C^23^ from cell populations at the cell cycle phases G1, S, and G2/M. We specifically included Bru-seq to capture nascent RNA transcripts for measuring functional output. We reason that instantaneous chromatin interactions between cis-regulatory elements (such as enhancers and promoters) determine gene transcription activities, therefore differential allelic expression could be best captured “live” by Bru-seq.

We developed an analytical framework that integrated haplotype-resolved Hi-C, RNA-seq, and Bru-seq data for fine dissection of the difference between the Pat and Mat genomes during the cell cycle. We studied the genome from a network point of view, where nodes of the network correspond to genomic loci partitioned at gene, topologically associating domains (TADs) defined by Dixon et al ^24^, and chromosome scales, and edge weights of the network correspond to contact numbers between two loci determined by Hi-C. This allowed us to extract multiple topological properties from the Hi-C data by using the concept of network centrality ^25^, which facilitate quantitative integration with RNA-seq and Bru-seq. The study of centrality, i.e., evaluating the degree of nodal importance to the network structure, can be used to identify and rank essential nodes (such as TADs and genes) in biological networks. A number of centrality measures exist. For example, degree centrality measures the total number of connections a node has, while closeness centrality measures the average distance of a given node to all other nodes. Eigenvector centrality assigns importance to a node based on the sum total of its neighbors’ importance,. These centrality measures extract important Hi-C features largely overlooked in previous studies, and provide an important way to understand the genome architecture and its dynamic process across cell cycle phases.

In this report, we discriminated the Pat and Mat genomes at the scales of whole chromosomes, TADs, inter-gene interactions, and intra-gene sequence variations in cis-regulatory elements. Using Bru-seq and RNA-seq, we found thousands of genes showing ABE with 98% of the dominant alleles expressed from one or the other parental origin throughout the cell cycle. We found homologous alleles switched between active and inactive chromatin compartment states, and the switched alleles often located close to compartment domain boundaries. We distinguish parental chromosomes with phase portraits ^26^ that provide a useful quantitative assessment of form-function dynamical relationships at the chromosome level. We also show that the TADs with the largest allelic differences are smaller in size and contain higher gene density compared to randomly selected TADs. Furthermore, we show a subset of ABE genes preferentially localized near TAD boundaries compared to random sampling of gene sets of the same size. We observed that ABE genes were enriched with SNVs/indels at CTCF binding sites. These results provide a comprehensive view of how the two haploid genomes differ in form and function, which may impact our understanding of human phenotypic traits and their penetrance, genetic diseases, vulnerability to complex disorders, and the development of cancers.

## Results

### Cell cycle-regulated gene expression

We analyzed bulk gene expression data from cell populations at G1, S, and G2/M sampled by Fluorescence-activated cell sorting (FACS) (Online Methods). From RNA-seq data sequenced on average 33 million reads per replicate, we identified 1451, 1388, and 1416 differentially expressed genes in G1 vs S, G2/M vs G1, and G2/M vs S pair-wise comparisons (false discovery rate, FDR < 0.05), respectively (Extended Data Figure 1A, Extended Data Table 1). When analyzing three replicas of Bru-seq data sequenced at a depth of ~40 million reads, we identified 568, 417, and 34 genes that were differentially transcribed in G1 vs S, G2/M vs G1, and G2/M vs S pair-wise comparisons FDR < 0.05) (Extended Data Table 2). As expected, Bru-seq showed for example that the G2 to M transition specific gene *CCNB1* was transcribed at a higher level in the G2/M population compared to that of G1 and S (Extended Data Figure 1B). We found 277, 235, and 21 differentially expressed genes common to both the RNA-seq and Bru-seq data sets in G1 vs S, G2/M vs G1, and G2/M vs S pare-wise comparisons (r= 0.92, 0.90, and 0.60), respectively (Extended Data Table 3), showing as expected a concordance between the rate of transcription and the steady-state levels of cytoplasmic mRNA during the cell cycle. Functional annotation showed that sets of genes identified from both Bru-seq and RNA-seq were significantly enriched under Gene Ontology (GO) terms related to the cell cycle (Extended Data Table 4).

### Allelically biased expression (ABE) during the cell cycle

We aimed to discriminate between the contribution of the Pat and Mat genomes to the dynamical functional human genome during the cell cycle. We first identified ABE genes from RNA-seq analysis. Specifically, from the 23277 genes interrogated, we identified 6795 transcripts containing informative heterozygous SNVs or indels. There were 5058 informative genes with allele read counts ≥5, which is the minimum number of counts that we used to reliably estimate ABE. Since the variables consisted of three cell cycle phases and two parental origins, we performed a two-way ANOVA analysis on the log_2_ transformed reads per kilobase per million reads (RPKM) values for the 5058 genes to identify ABE genes. This model identified 1762 (34.8%) genes (FDR < 0.05) that showed significant ABE (932 paternal allele high and 830 maternal allele high) (Extended Data Table 5). Of the 1762 genes, 713 were also differentially expressed when comparing between cell cycle phases G1, S, and G2/M (cell cycle-regulated). The set of ABE genes consists of 7.4% of the total 23277 genes interrogated, which is consistent with the fractions of ABE genes among human tissues studied ^18^. In terms of genome distribution, chr8 has the lowest number of ABE genes (5.45%), and chr22 has the highest percentage (11.17%)

For Bru-seq both exons and introns containing informative SNVs/indels were used to evaluate ABE. We identified 266,899 informative SNVs/indels from the Bru-seq data, while only 65,676 such SNVs /indels from RNA-seq data. However, in the Bru-seq data, many SNVs/indels had too low read depths (<5) to be included in the analysis. Using a read depth criterion ≥ 5, similar numbers of informative SNVs were found in the RNA-seq and Bru-seq data (19,394 and 19,998 respectively). Inclusion of the intronic SNVs increased the number of informative genes to 6,168. We observed relative larger variances between the three Bru-seq replicates compared to that of the RNA-seq replicates prompting us to apply the Tukey criteria ^27^ to select the top 5% genes with the largest allelic bias (Extended Data Table 6).

Among the ABE genes from RNA-seq or Bru-seq analysis, we found 103 common mono-allele expressed (MAE) genes (Extended Data Table 7). These include six known imprinted genes, four expressed from the Pat allele (*KCNQ1OT1, SNRPN, SNURF, and PEG10*) and one (*NLRP2*) from the Mat allele.

### Allele chromatin compartment switching between the Pat and Mat genomes

Cells were FACS-sorted into G1, S and G2-phases of the cell cycle and were then cross-linked with formaldehyde and subjected to Hi-C analysis (Online Methods). It is well known that interphase chromatin is partitioned into active euchromatin (A) and inactive heterochromatin (B) compartments ^23^. We constructed Hi-C maps at 100kb resolution for the Pat and Mat genomes. We also binned the RNA-seq and Bru-seq data into the same resolution of Hi-C binning. We observed that approximately 0.6% of 100kb bins switched compartment between the Pat and Mat alleles at each of the cell cycle phase G1, S, and G2/M (Extended Data Table 8), and the switched alleles were not the same in different cell cycle phase and had no concordance with ABE. These results were consistent with previous observations ^9^. Interestingly, we found that A/B compartment switching highly likely mapped to alleles adjacent to compartment boundaries where transition of active and inactive domains occurred ^23^ (Figure 1A, Extended Data Figure 2, Extended Data Table 8). We also discriminated the two genomes by allelic difference in gene expression (Bru-seq and RNA-seq) and in degree of Hi-C interactions (all expressed as log_2_ ratios between Pat and Mat) (Figure 1B). We noticed that A/B switching did not fully characterize Hi-C allelic difference, e.g., Hi-C degree difference (Figure 1C). When focused on the top 10% of bins with the largest Hi-C degree change, we observed that bins with significant Hi-C degree difference are associated with the significant differential allelic expression from Bru-seq analysis (P < 0.01), but not correlated with allelic difference from RNA-seq analysis (P = 0.2652). These results imply that the allelic difference of genome structure correlated well with ABE on nascent transcription as determined by Bru-seq analysis. Therefore, a form-function relationship is more accurately captured when comparing Hi-C data with nascent RNA Bru-seq data rather than with steady-state RNA-seq data.

**Figure 1.**
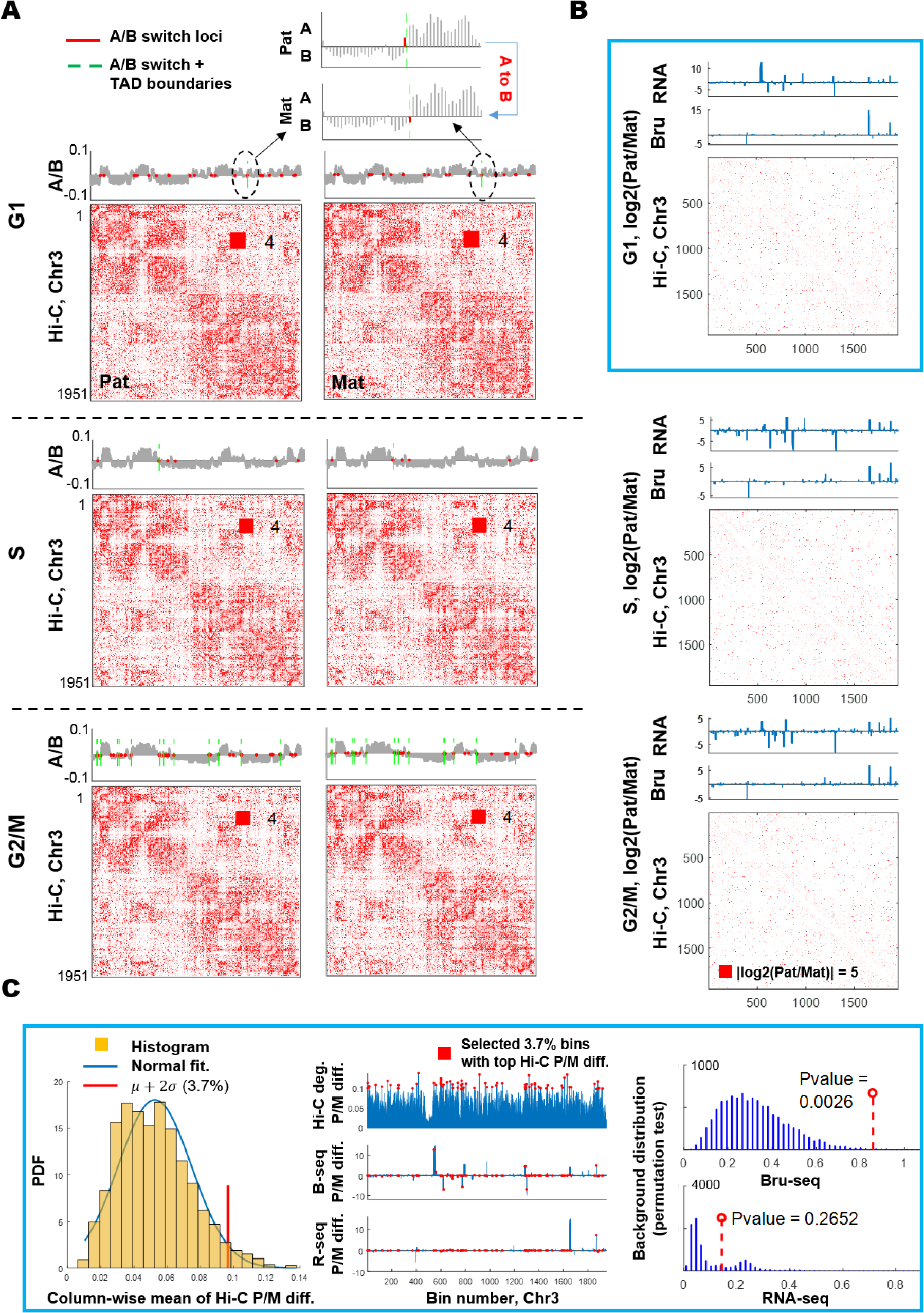
Identification of chromatin compartment switching regions and correlation of switched regions with gene expression change between the Pat and Mat genomes. A: Illustration of chromatin compartment switching between Pat and Mat on chr3. A/B compartments are plotted on top of each Hi-C map, with TAD boundary locations. A zoomed-in view of a switched region is shown on the top of the G1 Hi-C plot. The red square in Hi-C matrices represent the maximum number of contacts after applying the Hi-C normalization method ^26^. B: Characterization of form-function difference between homologous chr3. Here Pat or Mat represents the Pat-specific or Mat-specific Hi-C contact map and gene expression. In line plots, the X-axes of plots provide 100kb bin indices, and Y-axes correspond to log_2_(Pat/Mat) of RNA-seq or Bru-seq RPKM values, and in matrix plots, the X- and Y- axis are the genomic coordinates of chr3, and dot color represents the absolute value of log_2_(Pat/Mat) of normalized Hi-C contacts. C: Left shows the probability density function (PDF) of Hi-C contact allelic differences, given by the absolute values of log_2_(Pat/Mat), where Pat or Mat presents the column wise mean of Hi-C contact matrices for Pat allele or Mat allele. And the two times standard variation *(σ)* away from the mean (*μ*) of Hi-C contact allelic difference is the threshold to extract Hi-C bins (3.7%) with the largest allelic differences. Middle shows the locations of selected bins at chr3 (bar with read point head) and corresponding Bru-seq and RNA-seq difference between Pat and Mat. Right shows statistical significant test (Online Methods) of Hi-C difference correlation with gene expression differences in the same bins. Note that Bins with significant Hi-C magnitude difference yield significant Bru-seq allelic difference with P < 0.01.

### Chromosome phase portrait

To explore genome-wide differences between the Pat and Mat genomes, we introduce the concept chromosome phase portrait that provides a quantitative assessment of form-function dynamics at the chromosome level. This concept is in the same spirit of portrait of 4D Nucleome (4DN) introduced by Liu et al. ^28^. Specifically, we introduce a 3D space, where X and Y axes represent the genomic function in terms of RNA-seq and Bru-seq, and Z axis characterizes the genome structural property given by the network connectivity (also known as Fiedler number, FN) of the chromatin contact map ^26^. In general, the larger the value of FN is, the more well-organized a chromosome is (Online Methods). The proposed chromosome portrait allows us to describe each chromosome in a form-function domain, made up of two groups (Pat and Mat) at three cell cycle phases (G1, S, G2/M) (Figure 2A). As can be seen, chromosomes occupy distinct regions in the 3D space, and their paternal and maternal counterparts exhibit different form-function dynamics. Such an allelic difference becomes clearer when the chromosome portrait is projected to the RNA-seq – Bru-seq (X-Y) plane and the Bru-seq – FN (Y-Z) plane (Figure 2B). Note that the horizontal and vertical position shift in the X-Y plane reflects the change of allelically biased transcription in terms of RNA-seq and Bru-seq, respectively, while the vertical position shift in the Y-Z plane gives the structure change of two homologous chromosomes. In particular, we observed that the portrait of Pat chr9 consisting of G1, S, and G2/M is separated from that of the Mat counterpart, and the underlying dominant factor contributed to the allelic difference is gene expression in terms of RNA-seq. By contrast, the separation of Pat and Mat at chrX is largely contributed by Bru-seq. The form-function difference between every two homologous chromosomes is then summarized at each cell cycle phase (Figure 2C). Such a difference is characterized by the 2D distance of two allele-specific chromatin states in the proposed chromosome portrait. We found that chr9, 22 and X show larger differences than the other chromosomes. In addition to the allelic difference, the temporal form-function change along with the cell cycle can also be exhibited by measuring the difference between two consecutive time points (averaged over Pat and Mat homologs) (Figure 2C). We observed that evolution from G1 to S yields a larger form-function difference than that from S to G2/M. Taken together, using form and function information simultaneously improves discriminative power for a better understanding of allelic biases along with the cell cycle’s evolvement.

**Figure 2.**
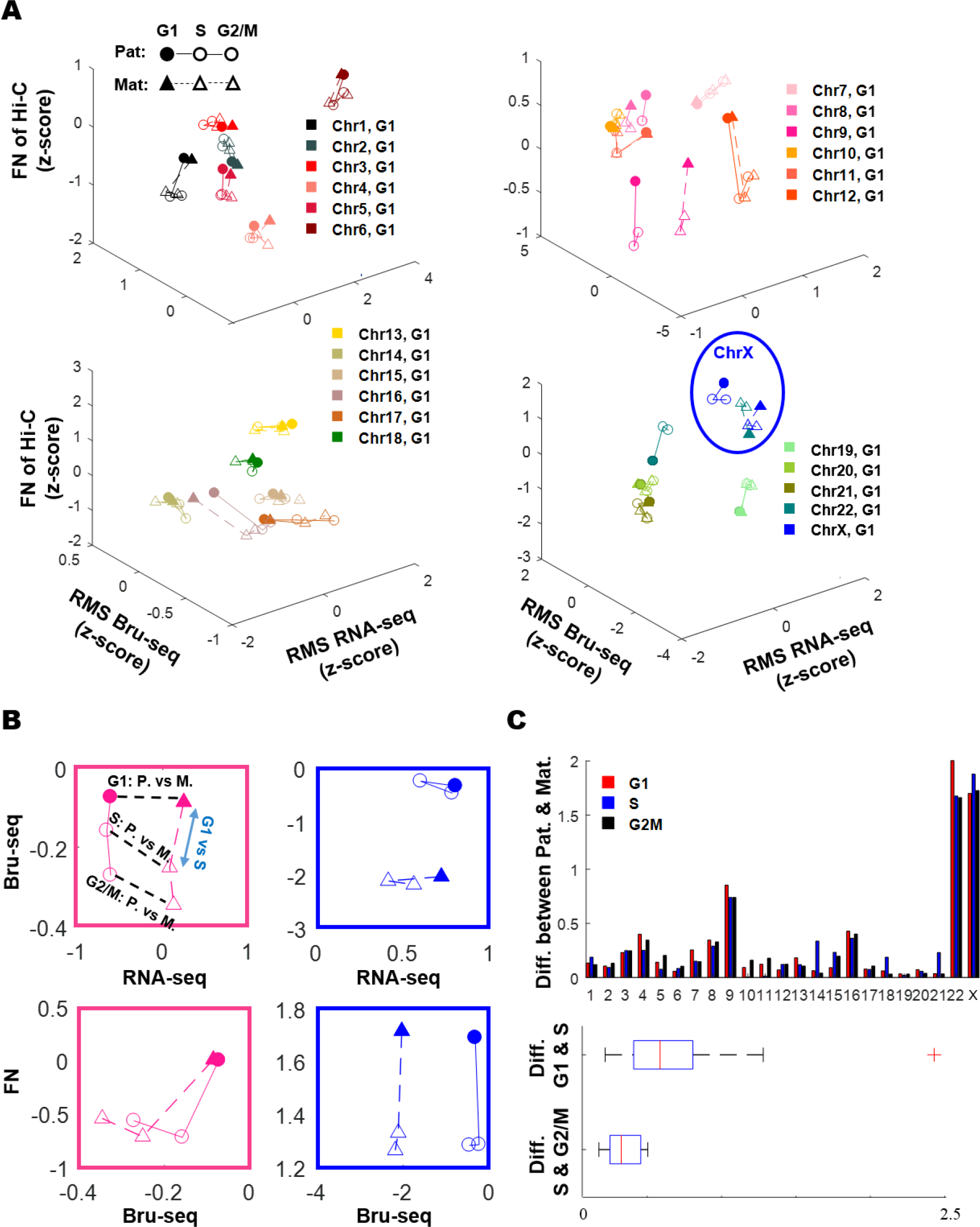
Phase portrait of chromosomes of the Pat and Mat genomes. A: Phase portrait of chromosomes at different cell cycle phases. Each chromosome is described by an allele-specific form-function domain (3D), made up of 3 cell cycle stages (G1, S, and G2/M) for the Pat and Mat homologs, respectively. We designate axis X as a measure of function given by z-scores of root mean square (RMS) of RNA-seq RPKM values, the axis Y as a measure of function in terms of z-scores of RMS of Bru-seq RPKM values, and the axis Z as a measure of form in terms of z-scores of the chromosome connectivity captured by Fiedler number. The filled square marker represents the cell cycle phase G1, and the other unfilled markers represent M and G2/M, respectively. The separation between the Pat portrait and the Mat counterpart implies the structural and functional difference. B: 2D projection of 3D phase portrait of Pat and Mat chr9 and X. Top: Allelically biased chromatin transcription on RNA-seq-Bru-seq plane. Bottom: Allelically biased chromatin form-function state on Bru-seq-FN plane. C: Form-function difference between the Pat and Mat homologs, given by their Euclidean distance in the phase portrait. D: Form-function difference between cell cycle stages: G1 to S, and S to G2/M, given by the Euclidean distance at two consecutive cell cycle stages averaged over Pat and Mat homologs for all chromosomes.

### Domain organization differences between the Pat and Mat genomes

In addition to A/B compartment partitioning, interphase chromatin is further organized into TADs that are conserved in vertebrates and are relatively cell-type invariant ^24^. We next explore allelic TADs difference between the Pat and Mat genomes. Previous work showed that the boundaries of TADs remained stable between cell types ^24^, and TADs domain organizations were relatively consistent between the Pat and Mat genomes ^9^. However, allelic biases at TAD-level interactions and functional changes at each cell cycle phase are not well understood. Our key idea is to interpret the genome as a network of TADs, where network vertices correspond to TADs whose loci are presented by *Dixon et al.*^24^, and edge weights are given by the contact frequency between two TADs from Hi-C. The function associated with a TAD is characterized by transcript abundance (in RNA-seq and Bru-seq RPKM values) of genes contained in it. Note that the perspective from network of TADs facilitates us to extract structural features of genome architecture, and the centrality analysis ^25^ (Online Methods) helps to identify TADs that play key topological roles in the allelically biased genome.

We applied the network centrality approach (Online Methods) and principal component analysis (PCA) to extract 2D representation of the allelically biased form-function features from our TAD-scale dataset (Figure 3A), in which each of the Pat and Mat genomes are mapped to a 2D point configuration at each cell cycle stage. We found that eigenvector centrality, betweenness centrality, degree centrality, and local Fiedler vector centrality (LFVC) are the top network centrality features (see details on centrality measures in Online Methods) to discriminate the Pat and Mat genomes at TAD scale. We also noted that although the pattern of the genome in terms of TAD loci plotted in the 2D space is similar between the Pat and Mat genome in general (Figure 3A), positions of allele-biased TADs apparently shifted, e.g., TAD at Chr22 from bin 392 to bin 408 at 100Kb unit (Figure 3A). This implies that there exist some TADs that delineate allelic chromatin modifications and transcription. Spurred by that, we measured the distances between the Pat TAD and the Mat TAD (Figure 3B). Focusing on the top 10% TADs whose allele-specific positions change the most (Extended Data Table 9), we found that chr9, 22 and X contain TADs with extremely large Pat-Mat difference. This result is consistent with the allelic bias identified by chromosome portrait shown in Figure 2C. We further investigated the structure and transcription properties of the selected TADs. As can be see, these TADs were significantly different in allelic expression and degree centrality of Hi-C, and interestingly, they had higher gene density and were smaller in size than randomly selected TADs (Figure 3B right); statistical significance is found by using a randomization test (Online Methods). The above results indicate that our identified allelically biased TADs may represent form-function deviations between the Pat and Mat genomes.

**Figure 3.**
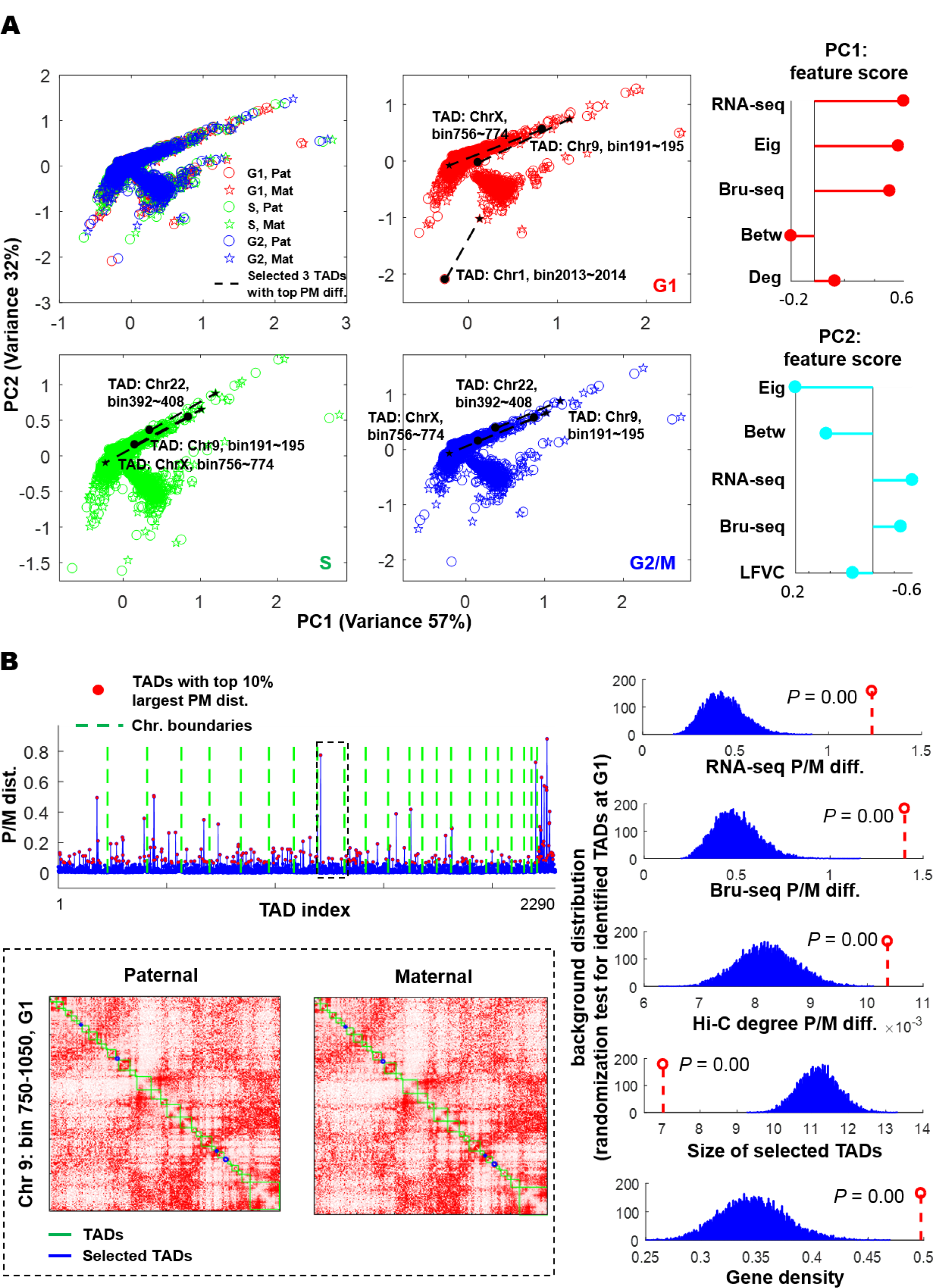
Allelically biased form-function dynamics of TADs. A: 2D representations (PCA) of form-function features of allelically biased TADs at different cell cycle stages. Left: Principal component (PC) analysis of form-function features of 2290 TADs in the whole genome. The circle (or star) marker represents a TAD corresponding to the Pat (or Mat) genome. At each cell cycle stage, TADs with the top three largest Pat-Mat distance are marked. Right: Contribution of function-form features on each of PCs. In PC1 and PC2, RNA-seq and Bru-seq describe the function of allele-specific genome, eigenvector centrality, betweenness centrality, degree centrality and local Fiedler vector centrality (LFVC) to characterize the topological properties of Hi-C contact maps at TAD scale. A feature score with larger magnitude implies a larger contribution of the corresponding feature to PCA. B: Characteristics of TADs with top 10% paternal-maternal difference. Left-top: Pat-Mat difference versus TAD index, where the former is quantified by the Euclidean distance of allele-specific TADs on the PC1-PC2 plane. Left-bottom: Example of selected TADs on chr9. Right: Statistical significance of form-function differences (top three plots) and genome characteristics (TAD size and gene density in bottom two plots) of the selected TADs.

### Seeing allelic biases from multilayer inter-gene contact networks

Besides the TAD-scale analysis, one may ask the questions whether there exist inter-gene (namely, gene-to-gene) contact differences between the Pat and Mat alleles, and how such differences if any evolve through the cell cycle phases. To incorporate the impact of cell cycle phases, we interpret the allele-specific inter-gene contact maps as a multilayer network ^29^, in which each layer corresponds to a cell cycle phase that associates with an inter-gene contact network that is generated from Hi-C data. In our multilayer network analysis, we focused on the set of 564 genes that showed ABE from Bru-seq analysis, and were also assessed for ABE in RNA-seq analysis (Extended Data Table 5). We determined the Pat/Mat inter-gene network based on raw Hi-C contacts, where there existed an edge between two genes if the corresponding contact number is nonzero (Figure 4A). As can be seen, the number of intergene contacts varied between the Pat and Mat alleles, leading to different network topologies that show dynamical changes during the cell cycle (Extended Data Figure 5). To distinguish the Pat allele from the Mat allele, we adopted the overlapping degree centrality and the multiplex participation coefficient to quantify the distribution of the node degree across cell cycle phases (Figure 4B and Online Methods). Here the former metric identifies hubs from networks, and the latter quantifies the participation of a gene to the different network layers, namely, cell cycle phases. With respect to the Z-score of genes’ overlapping degree *O*, we distinguished hubs (involving a large number of interactions), for which *O* ≥ 2, from regular nodes (non-hubs), for which *O* < 2. With respect to the multiplex participation coefficient *M*, we called focused genes if their degrees were concentrated at a single layer, corresponding to 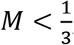, and multiplex genes if their connected edges were homogeneously distributed across the three layers, corresponding to 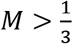. We found that there exist genes having largely varied topological roles on Pat and Mat alleles. Spurred by that, we extracted the top 10% genes whose positions change the most due to allelic biases in the 2D plane formed by overlapping degree and multiplex participation coefficient (Figure 4C and Extended Data Table 10). Our analysis revealed two classes of inter-gene contacts: intrachromosome contact (the interacted genes belong to the same chromosome), and inter-chromosome contact (the interacted genes fall into different chromosomes). As we can see, four possible scenarios existed: a) genes with more intra-chromosome contacts on the Mat homolog (e.g., SPTBN1), b) genes with more inter-chromosome contacts at the Mat homolog (e.g., NCALD), c) genes with more intrachromosome contacts at the Pat homolog (e.g., STK17B), and d) genes with more inter-chromosome contacts at the Pat homolog (e.g., LEPREL1). The above results imply that the inter-gene contact differences reflect allelic chromatin modifications that evolve through cell cycle phases, and further suggest that the networked ABE genes may be transcriptionally co-regulated or co-transcribed in transcription factories.

**Figure 4.**
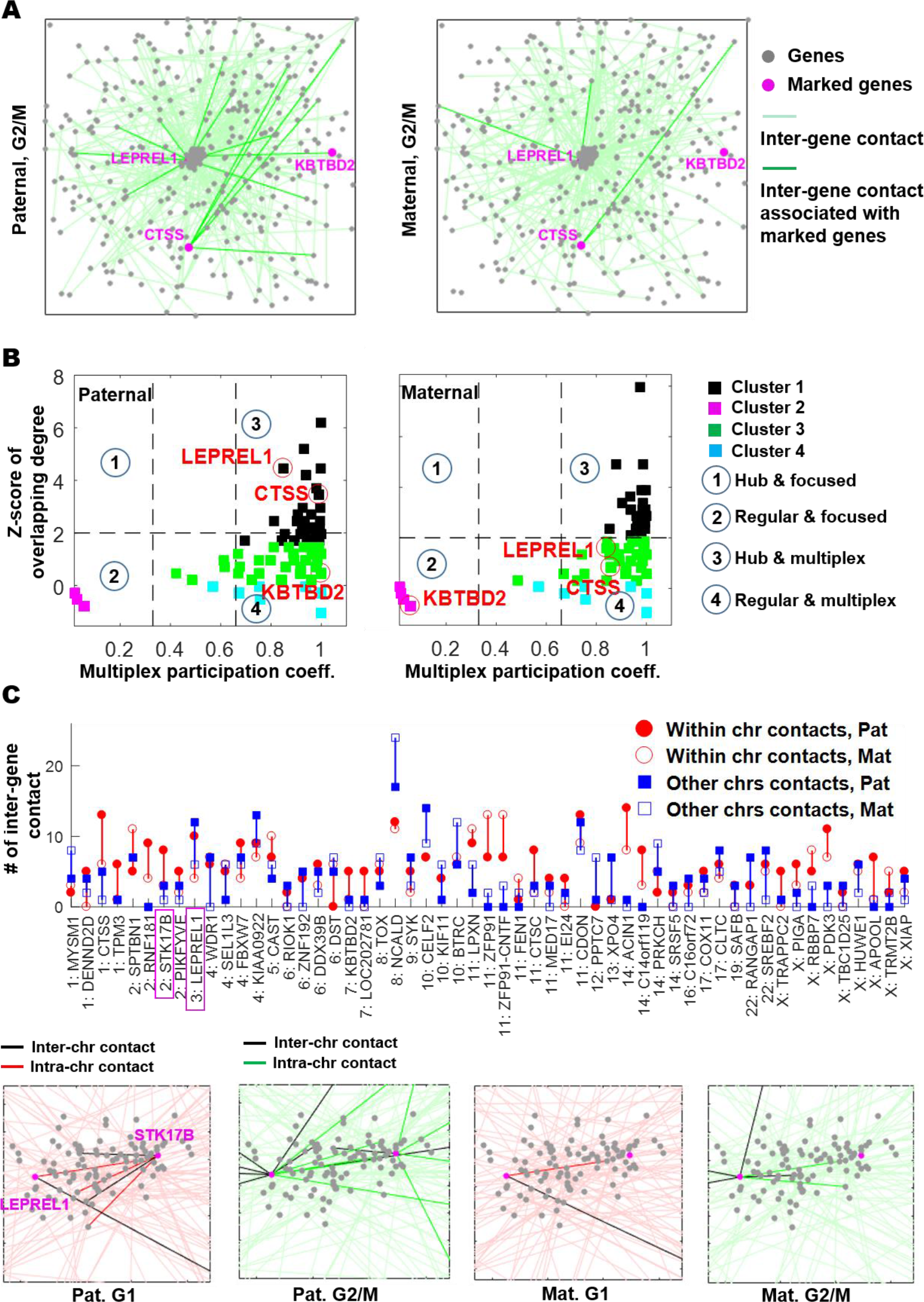
Evolution of allele-specific inter-gene contact network over cell cycle phases. A: A representative allele-specific inter-gene network at G2/M, where nodes correspond to ABE genes (See Extended Data Table 5), and edges represent the presence of contacts between two genes. The positions of nodes are determined by a layout engine in MATLAB (*dolayout*) for a bio-graph object (*biograph*) generated by gene-level Hi-C. B: Characteristics of allelically biased inter-gene contact network, which include the overlapping degree (O) and the multiplex participation coefficient (M) (Online Methods). The larger the value of O, the more contacts a gene has during all cell cycle stages. Furthermore, the larger the value of M, the more equally distributed is the participation of a gene to all cell cycle stages. In the O-M plane, we divide genes into 4 clusters by using the K-means algorithm. C: Intra-and inter-chromosome contacts of the top 10% genes with the largest paternal-maternal difference (ranked by their position shift in the O-M plane). Top: Summary of allele-specific intra-and inter-chromosome contacts of the selected genes. Bottom: Evolution of allele-specific inter-gene contact for gene STK17B and LEPREL1.

### Cis-regulatory elements (cis-REs) variations in ABE genes

We also evaluated a set of 216 significant ABE genes common to Bru-seq and RNA-seq for their allelic intra-gene form-function relationship. We found that 72 of these ABE genes were localized in TAD boundary regions, which include TAD boundaries at 100Kb bin unit and their adjacent bins at both sides. This result is statistically significant when compared to a random sampling of the same number of non-ABE genes (Figure 5A, Extended Data Table 11). To gain insights into the underlying mechanism for the biological meaning of our observed results and understand why many ABE genes are localized on TAD boundary regions, we focus on analyzing cis-regulatory elements (cis-REs), including known enhancers, promoters, the CCCTC transcription factor (CTCF) binding sites, RAD21 (one of the subunit of cohesin) binding sites and SNV/indels around and within the genes. We observed that there were no significant differences in the densities of promoters, CTCF sites, RAD21 sites, and SNVs/indels in the genomic regions of the gene analyzed compared to randomly sampled genes of the same size. The one variance, the number of SNVs/indels in CTCF binding regions, is statistically different compared to random sets of genes size matched (Figure 5B left), which provide further evidence to support that CTCF is the key organizer for chromatin architecture ^30,31^. Furthermore, we derived the first two principal components of genes’ cis-regulatory elements, and identified a subset of genes with largely distinct cis-regulatory element features (Figure 5B right). We then generated intra-gene Hi-C maps (at 5kb resolution) given the genomic coordinates to include the gene body plus 25 kb and 5 kb sequences to the 5’ and 3’ end of each gene. We found that a significant portion of ABE genes showed both form-function differences between the Pat and Mat alleles. It is clear from the identified gene *BAGALT3* (Figure 5C) that the local (intra-gene) allelic organization contributes to allelically biased expression. This reveals the unique roles of gene organization and allele preference for expression.

**Figure 5.**
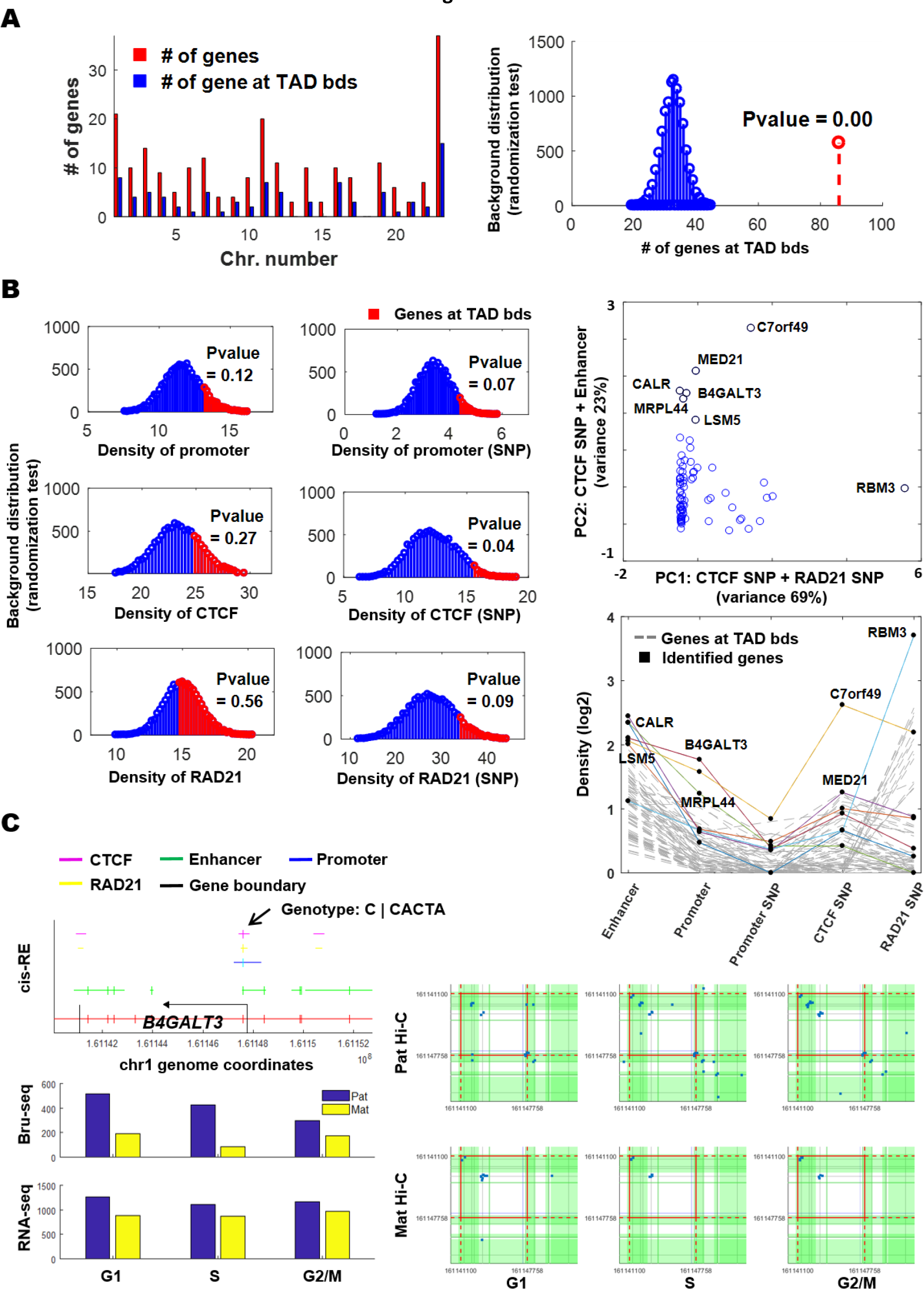
Characterization of intra-gene features for selected ABE genes. A: Chromosome distribution of the selected ABE genes (left), and a large portion of genes tend to localize on TAD boundary regions (right). Those regions contain TAD boundaries with +/− 1 bin (100 kb resolution) away from them. B: Comparison of Cis-REs (i.e., promoter, enhancer, CFCT and RAD21 binding site) and SNV/indel distribution in these elements for genes on TAD boundary regions against randomly picked genes (left); Identification of a subset of genes distinct from other genes by using PCA and Kmeans (right top), and genes with higher density of cis-regulatory elements or containing more SNVs/indels are plotted and identified by solid lines and points (right bottom). C: A composite view of the ABE gene *B4GALT3*. On the top left panel, genomic regions containing enhancers, promoter, CTCF sites, and RAD21 sites (cis-regulatory element, cis-RE) are indicated with color-coded lines; SNV/indel sites are shown with vertical bars; pointed by an arrow is genotype of an indel common to the promoter, an enhancer, and a CTCF/RAD21 binding region. Left bottom two panels show RPKM values of the Pat and Mat alleles measure by Bru-seq and RNA-seq at G1, S, and G2/M. The Hi-C interactions formed around this indel region disappeared in the Mat genome (right bottom 3 panels) compared to the Pat genome (right top 3 panels), the red box denote the genomic coordinates of *B4GALT3*, blue dots are Hi-C paired-end reads (interactions), and green shaded regions highlight predicted enhancers.

We also searched for enrichment of other transcription factor binding sites in ABE genes mapped to TAD boundaries. We identified 36 transcription factors (Extended Data Table 12) from a union of public databases that were shared among these genes. Extended data Figure 6 shows that pathway analysis and network-based gene set enrichment analysis generated using Ingenuity Pathway Analysis (IPA) ^32^. The consensus set of transcription factors bound to these ABE boundary genes were a subset of larger network that was significantly associated with “chromatin organization during cell development” at 1.29E-58 and “hormone regulation of the cell cycle” at 2.45E-23. Other members of the network include XIST (X Inactive Specific Transcript, ESR1 (estrogen receptor), AR (androgen receptor) and core histone complexes H3 and H4. These results confirmed that these transcription factors were involved not only in the regulation of gene transcription, but also in hormone-mediated chromatin re-organization during the cell cycle. This suggests that the sequence variations might explain a mechanism for ABE.

## Discussions

We report here on an integrated approach for the analysis of Hi-C, Bru-seq, and RNA-seq data for a human lymphoblastoid cell line, NA12878, whose diploid genome has been determined ^33,34^. We segregated its nucleome into the Pat and the Mat components distinguished by chromosome conformation and ABE at the cell phases G1, S, and G2/M determined by Hi-C, nascent RNA Bru-seq and steady-state RNA-seq. We showed that the difference between each parental homolog could be identified at the chromosome level. We found that at each cell cycle phase examined with Hi-C, switches between the euchromatin and heterochromatin compartment between the two genomes tended to occur at domain boundaries; ABE genes preferentially localized at TAD boundaries; allelic differences occurred in smaller sized TADs and there was a higher gene density in these TADs. We also identified differential inter-gene networks between the Pat and Mat loci from the analysis of form-function relationship between ABE genes, which might suggest that networked genes expressed from the same parental alleles transcriptionally co-regulated. Furthermore, we observed that the number of CTCF sites in ABE genes contained a higher number of sequence variations (e.g., SNVs and indels). Our results extend previous findings that sequence variations form the basis for allelically biased transcription factor binding, which in turn governs chromatin architecture and expression with allelic preference ^9^.

It is known that chromatin compartmentalization into euchromatin (A) and heterochromatin (B) domains correlates to transcriptional active and inactive regions ^23^, respectively. Here we discovered that A/B compartment switching occurs at each cell cycle phase, and the switched alleles were not the same from phase to phase during the cell cycle (Extended Data Table 8). This might be an expected phenomenon since the organization of the genome is dynamical during the cell cycle in proliferating cells. On the other hand, it is possible that the cells might be a collection from a wide range of unsynchronized cells sorted into corresponding cell cycle phases, especially in S phase the cells were a mixture from early stage to completed replication, and also in the long G1 phase. To exclude this possibility, it may require further analysis on similar data obtained from cells synchronized at specific time points during the cell cycle. One feature of our findings is that there is concordance between the difference of Hi-C interaction degree and ABE measured by Bru-seq (but RNA-seq). This observation suggests that interactions between cis-regulatory elements that drive gene expression ^35^ are instantaneous events, and such form-function relationship can be better captured by measuring nascent RNA rather than steady-state RNA. We also introduced a chromosome characterization method called phase portrait. This procedure takes form and function of each chromosome as a whole into account, so that the dominant component(s) from Bru-seq, RNA-seq, and Hi-C are easily identified and segregates the parental homologues in a 3D portrait space ^28^. This provides a useful tool for identifying chromosomes with distinct phase portrait space distribution related to form-function changes, either physiologically or pathologically.

As demonstrated by our bin-level, TAD-level, and gene-level analyses, network science has emerged as a powerful conceptual paradigm in biological science. However, little effort has been involved in nucleome dynamics under an integration of form and function. With the aid of network-based approaches such as graph theory, network centrality, and multilayer network theory, we are able to study genome structure from multiple views, and facilitate quantitative integration with functional information. The detailed connections between network structure and function in the context of the nucleome can be further studied in conjunction with novel experimental approaches, and can shed light on the importance of considering genetic variants in understanding haplotype-resolved genomics. A deeper understanding of nucleome form-function relationship will may have broad translational impact spanning cancer cell biology, complex disorder development, and precision medicine.

### Online Methods

We grew the NA12878 cells in RPMI1640 medium supplemented with 10% fetal bovine serum (FBS). Live cells were stained with Hoechst 33342 (Cat # B2261, Sigma-Aldrich), and then subjected to Fluorescence-activated cell sorting (FACS) to obtain cell fractions at the corresponding cell cycle phases G1, S, and G2/M. Cells used for Hi-C libraries construction, we cross-linked the cells with 1% formaldehyde after staining with Hoechst, neutralized the cross-linking reaction with 0.125M of glycine, and the cell were then subjected to FACS. Cells used for Hi-C, RNA-seq, and Bru-seq analyses were sorted live just after Hoechst staining.

### Hi-C library construction

Hi-C library construction and Illumina sequencing using established methods ^8^. Briefly, Cross-linked chromatin was digested with the restriction enzyme MboI for 12 hours. The restriction enzyme fragment ends were tagged with biotin-dATP and ligated in situ. After ligation, the chromatins were de-cross-linked, and DNA was isolated for fragmentation. DNA fragments tagged by biotin-dATP in the size range of 300–500 bp were pulled down for sequencing adaptor ligation and polymerase chain reaction (PCR). The PCR products were then sequenced on the Illumina HiSeq2500 platform in the University of Michigan DNA Sequencing Core facilities (Seq-Core).

### Alignment to NA12878 diploid genome

NA12878 diploid genome reference data was downloaded from AlleleSeq (Rozowsky, Abyzov et al. 2011) server. Trimmomatic (Bolger, Lohse et al. 2014) was used to trim Illumina adapter sequences and low quality ends from raw reads. Trimmed reads were then aligned using Bowtie2 (Langmead and Salzberg 2012) against paternal and maternal genomes separately, with R1 reads and R2 reads aligned separately. The paternal and maternal alignments were processed using the program “mergeBowtie.py” from AlleleSeq package. This program filters the alignments according to these criteria: (1) if a read aligns to one parental genome with fewer mismatches than the other, it is assigned to this parental genome only; (2) if a read aligns to both genomes at the same position with the same number of mismatches, it is kept for both; and (3) if a read aligns to both genomes with the same number of mismatches, but at different positions, it is discarded. The filtered paternal and maternal alignments were converted to BED format and then converted to hg19 coordinates using CrossMap (Zhao, Sun et al. 2014). The hg19-to-paternal/maternal chain files were obtained from the diploid genome reference data package and swapped to the other direction using program chainSwap downloaded from UCSC Genome Browser Utilities (Kent, Sugnet et al. 2002). BEDTools (Quinlan and Hall 2010) was used to intersect the alignments with all heterozygous variant positions in NA12878 obtained from the diploid genome reference package. Alignments were selected so that (1) either this read or its mate intersected with a heterozygous variant, and (2) both this read and its mate were assigned to the same parental genome. The alignments with mapping quality no lower than 20 were selected for Hi-C analysis.

We generated Hi-C data for populations at cell cycle phases G1, S, and G2/M for NA12878 cells. Respective to G1, S, and G2/M, we obtained 512.735, 550.288, and 615.226 million raw Hi-C sequence reads. On average, 62% of the raw reads were uniquely mapped to the genome of hg19 assembly. Of the mapped, on average we are able to identify 4.339 million paired-end reads containing heterozygous SNPs or short sequence variants. We used these informative reads to construct the diploid Hi-C matrices, consisting of haplotype-resolved Hi-C maps for the paternal Pat and maternal Mat genomes at G1, S, and G2/M. All Hi-C matrices were generated using the software HOMER ^36^.

### Determine mRNA abundance using RNA-seq analysis at G1, S, and G2/M

The live cells sorted were used for total RNA extraction. Subsequently RNA-seq library construction was carried out in the seq-core facility, and sequence reads of 50-base in length were generated on an Illumina HiSeq 2500 station.

### Determine allele-specific nascent transcription using Bru-seq analysis at G1, S, and G2/M

We performed 5’-bromouridine (Bru) incorporation in live cells for 30 minutes, and the Bru-labeled cells were then stained on ice with Hoechst 33342 for 30 min before subjected to FACS at 4°C to isolate G1, S, and G2/M phase cells. The sorted cells were immediately lysed in TRizol (Cat # 15596026, ThermoFisher) and frozen. To isolate Bru-labeled RNA, DNAse-treated total RNA was incubated with anti-BrdU antibodies conjugated to magnetic beads ^21^. We converted the Bru-labeled transcripts from the different samples into cDNA libraries and they were deep-sequenced at 50-base length on an Illumina HiSeq2500 platform.

### Allele specific RNA-seq and Bru-seq analyses

We developed a pipeline for ABE analysis. The pipeline developed for estimating allele specific expression from RNA-seq and Bru-seq is outlined in Figure 3. The left side shows the normal flow of a non-allele specific RNA-seq or Bru-seq pipeline which is combined at the end with results from the allele specific portion of the pipeline to get abundance estimates for the maternal and paternal copy of each gene.

The normal RNA-seq and Bru-seq analysis were performed as previously described (Seaman 2017 and Paulsen 2013, respectively). Briefly, Bru-seq used Tophat (v1.3.2) to align reads without de novo splice junction calling after checking quality with FastQC. A custom gene annotation file was used in which introns are included but preference to overlapping genes is given on the basis of exon locations and stranding where possible (See Paulsen 2013 for full details). Similarly, in RNA-seq data process, the raw reads were checked with FastQC (version 0.10.1). Tophat (version 2.0.11) and Bowtie (version 2.1.0.0) were used to align the reads to the reference transcriptome (HG19). Cufflinks/Cuffdiff (version 2.2.1) was used for expression quantification and differential expression analysis, using UCSC hg19.fa and hg19.gtf as the reference genome and transcriptome. A locally developed R script using CummeRbund was used to format the Cufflinks output.

To determine allele specific transcription and gene expression through Bru-seq and RNA-seq, all reads were aligned using GSNAP, a SNV aware aligner (Wu 2010). Hg19 and USCS gene annotations were used for the reference genome and gene annotation, respectively. The gene annotations were used to create the files for mapping to splice sites (used with −s option). Optional inputs to perform SNV aware alignment were included. Specifically, −v was used to include the list of heterozygous SNVs (ftp://platgene_ro@ussd-ftp.illumina.com/2016-1.0/hg19/small_variants/NA12878/NA12878.vcf.gz) and --use-sarray=0 was used to prevent bias against non-reference alleles.

After alignment, the output SAM files were converted to BAM files, sorted and indexed using SAMTOOLs (Li 2009). SNV alleles were then quantified using bam-readcounter (D. Larson et al., https://github.com/genome/bam-readcount) which was used to count the number of each base that was observed at each of the heterozygous SNV locations. The statistical significance of allele specific expression for each SNV was then quantified by a binomial test with a null probability of 0.5.

Allele specificity of each gene was then assessed by combining all of the SNVs in each gene. For RNA-seq only exonic SNVs. For Bru-seq exonic SNVs were counted first, and then non-exonic SNVs were counted if they were in a gene’s intronic region. Paternal and maternal abundance of each gene were calculated by multiplying the overall abundance estimate by the fraction of the SNV-covering reads that were paternal and maternal, respectively.

Gene level significance of allele specific expression for each gene in each cell cycle stage was evaluated using a negative binomial model variance estimation is improved through a local regression relating variance to the mean (https://www.mathworks.com/help/bioinfo/ref/nbintest.html). The same method was used to determine differential gene expression between the overall abundance in different cell cycle stages. ANOVA was used on the log_2_ RPKM values to determine what genes changed over the cell cycles as well as between maternal and paternal alleles.

RNA-seq and Bru-seq were binned into 100kb and 1 Mb bins to match the resolution of the Hi-C data. This was done separately using the maternal and paternal expression estimates by adding the expression of the genes in a bin and when necessary dividing a gene’s counts according the proportion of the bin in each gene. About 75% of genes could not be assessed for allele specific expression due to a lack of SNVs in the gene body. When binning RNA-seq and Bru-seq, an assumption of 50% maternal and paternal expression was made for these genes to avoid losing that data.

From the 23277 Refseq genes interrogated, we identified 6795 transcripts containing informative heterozygous SNVs or indels. There were 5058 informative genes with allele read counts ≥5, which is the minimum number of counts that we use to reliably estimate ABE. Since the variables consisting of three cell cycle phases and two parental origins, we performed two-way ANOVA on the log_2_ transformed RPKM values for the 5080 genes to identify ABE genes. This model identified 1762 (34.8% of informative) genes (FDR < 0.05) that showed significant ABE (932 paternal allele high and 830 maternal allele high). Of the 1762 genes, 713 were also cell cycle regulated. In terms of genome distribution, on average there were 7.34% ABE genes among the chromosomes, chr8 has the lowest number of ABE genes (5.45%), and chr22 has the highest percentage (11.17%). The set of ABE genes consists of 7.4% of the total 23277 genes interrogates genes in our current work, which is consistent with the fractions of ABE genes among human tissues studied ^18^.

### Identification of ABE genes from Bru-seq analysis

Bru-seq detects nascent transcripts containing both exons and introns. Sequence reads from both exons and introns containing informative SNVs are used to evaluate ABE. We identified 266,899 informative SNVs from the Bru-seq data, while only 65,676 such SNVs from RNA-seq data. However, in the Bru-seq data, many SNVs show low read coverage depth to be able to be statistically evaluated compared to that in the RNA-seq data. We require allelic count ≥ 5 to estimate ABE for nascent transcripts as done for the RNA-seq data. This criterion found that there were similar numbers of informative SNVs (19,394 and 19,998) in the RNA-seq and Bru-seq data, respectively. Inclusion of the intron SNVS makes the number of informative genes to 6,168 for ABE estimation.

We observed that there were relative large variances between samples in the Bru-seq data set. Due to the increased variability between replicates in the Bru-seq data compared to the RNA-seq data, instead of using two-ANOVA, we used another method to identify ABE genes. In which we subtracted the expression values of each gene in the maternal samples from those in the paternal samples. We then identified 563 genes with ABE in G1, 594 in S, and 610 in G2/M, which yield the top 5% largest differences in allelic expression. There were 1039 unique genes from G1, S, and G2/M combined.

### Transcription factor binding sites analysis

Transcription factor binding to boundary genes was defined using the union of data from public data sources, including the FactorBook Motif pipeline ^37^, ENCODE project consortium ^33^ and the NIH roadmap epigenomics mapping consortium ^38^. Gene set enrichment analysis was performed using Panther Amigo2 gene ontology ^39^ and network-based enrichment using Ingenuity Pathway Analysis^®^ (Qiagen GmBH) ^32^.

### Statistical significance via randomization test

A randomization test builds the shape of null hypothesis (namely, the random background distribution) by resampling the observed data. In our analysis, unless specified otherwise, this sampling procedure was repeated 1000 times, and a rank-based P value is then calculated for the right or left-tailed event. For example, the background distribution in in Figure 2C is generated by calculating the average gene expression for randomly selected Hi-C bins. And the probability of the right-tailed event at our observation under the background distribution yields the P value. Similar statistical tests were used in Figure 4 and 6.

### Network connectivity measure: Fiedler number

We regard a Hi-C contact map as a graph, where graph vertices correspond to biological units, such as bins, TADs, and genes, and edge weights are given by the interaction frequency between two vertices. More formally, let *G* = (*V*, *E*) represent an undirected graph where *V* is the set of nodes with cardinality |*V*| = *N*, and *E* ⊆ {1,2,…,*N*} × {1,2,…,*N*} denotes an edge set. The Hi-C matrix ***H*** can be interpreted as an adjacency matrix of *G* by removing its diagonal elements. And the corresponding graph Laplacian matrix is given by ***L*** = ***D*** − ***H***, or its normalized version 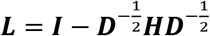, where ***D*** is the degree matrix ***D*** = *diag*(***H*1**), **1** is the vector of all ones, and *diag*(***x***) signifies the diagonal matrix with the diagonal vector ***x***. It is known from spectral graph theory ^40^ that Fiedler number of *G* is given by the second smallest eigenvalue of the graph Laplacian ***L***. The magnitude of this value reflects how well connected the overall network is.

### Network centrality analysis

A network/graph centrality measure is a quantity that evaluates the influence of each node to the network, and thus provide essential topological characteristics of nodes ^25^ (Extended Data Figure 3). In this paper, we extract structural features from Hi-C contact maps by using centrality measures discussed in ^28^: degree centrality, eigenvector centrality, local Fiedler vector centrality (LFVC), closeness, betweenness, local clustering coefficient (LCC), weight of multi-hop walks, and distance to reference nodes. Here a nodal degree is defined as the sum of edge weights (namely, Hi-C contacts) associated with each node. The eigenvector centrality is defined as the principal eigenvector of the adjacency matrix corresponding to its largest eigenvalue, which measures a node’s influence to the entire network based on its neighbors’ influence. LFVC evaluates the structural importance of a node regarding the network connectivity. Closeness is defined by the shortest-path distance of a node to all other nodes, which implies how far one node is from the geometrical center of a network. Betweenness is the fraction of the number of shortest paths passing through a node relative to the total number of shortest paths in a connected network. A node with high betweenness has the potential to disconnect the network if it is removed. LCC of a node quantifies how close its neighbors are to being a complete graph. The h-hop walk weight of a node is given by the sum of edge weights associated with paths departing from this node and traversing through h edges. Given nodes of interest, we can explore network distances of each node to the reference nodes as structural features. The use of network distances helps to avoid ambiguity of centrality measures due to the possibly high structural symmetry in networks.

### Overlapping degree and multiplex participation coefficient in multilayer networks

In our proposed multilayer network, genes (namely, ‘nodes’) are connected by both intra-layer and inter-layer connections (namely, ‘edges’), where the intra-layer connection corresponds to the contact frequency between genes, and the inter-layer connection gives a visual link between a gene and its counterpart at two cell cycle phases (Extended Data Figure 4). The overlapping degree and the multiplex participation coefficient are introduced to study how the nodal degree is distributed in a multilayer network. Let *M* denote a multilayer network with *N* nodes and *L* layers, the degree of node *i* on layer *α* is given by 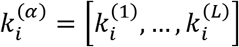. The overlapping degree of node *i* is then given by 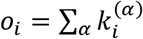. It can be used to identify hubs from the network. However, one node that is a hub in one layer may only have few connections in another layer. Therefore, it is desired to quantify the participation of a node to the various layers. A suitable quantity to describe the distribution of edges connected to node *i* is the multiplex participation coefficient ^41^. 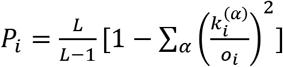. Here *P*_*i*_ takes values in [0,1] and measures whether or not the degree of node *i* is uniformly distributed among the *L* layers. If *P*_*i*_ = 1, then node *i* has exactly the same number of edges on each layer. If *P*_*i*_ = 0, all the edges of node *i* are concentrated in just one layer. The multiplex participation coefficient helps to capture the nodal dynamics in multilayer networks.

## Acknowledgements

We thank the University of Michigan sequencing Core member for producing high quality Bru-seq, RNA-seq, and Hi-C data. We thank Michele Paulson for generating the Bru-seq sequencing libraries. We also thank Dr. Jacob Kitzman for sharing the NA12878 cell line and assisting in resolving genotype-phasing issues of the cell line. This work is funded by a grant from the Forbs Foundation, and is supported, in part, by the DARPA Biochronicity Program and the DARPA Deep-Purple and FunCC Program.

## Author contributions

I. R. conceived and supervised the study. H. C. designed and performed the experiments. S. L. and L. S. performed computational analyses and interpreted the data. H.C. and M.L. performed RNA-seq and Bru-seq bulk data analyses, respectively. C.N. and H.C. carried out cis-regulatory element analysis. W.W. generated haplotype-resolved Hi-C matrices. G.H. performed transcription factor binding site enrichment and pathway analyses. All authors participated in the discussion of the results. S. L. and H. C. prepared the manuscript with input from all authors.

## Materials & Correspondence

Correspondence and material request should be addressed to IR.

## Extended Data

**Extended Data Figure 1.**
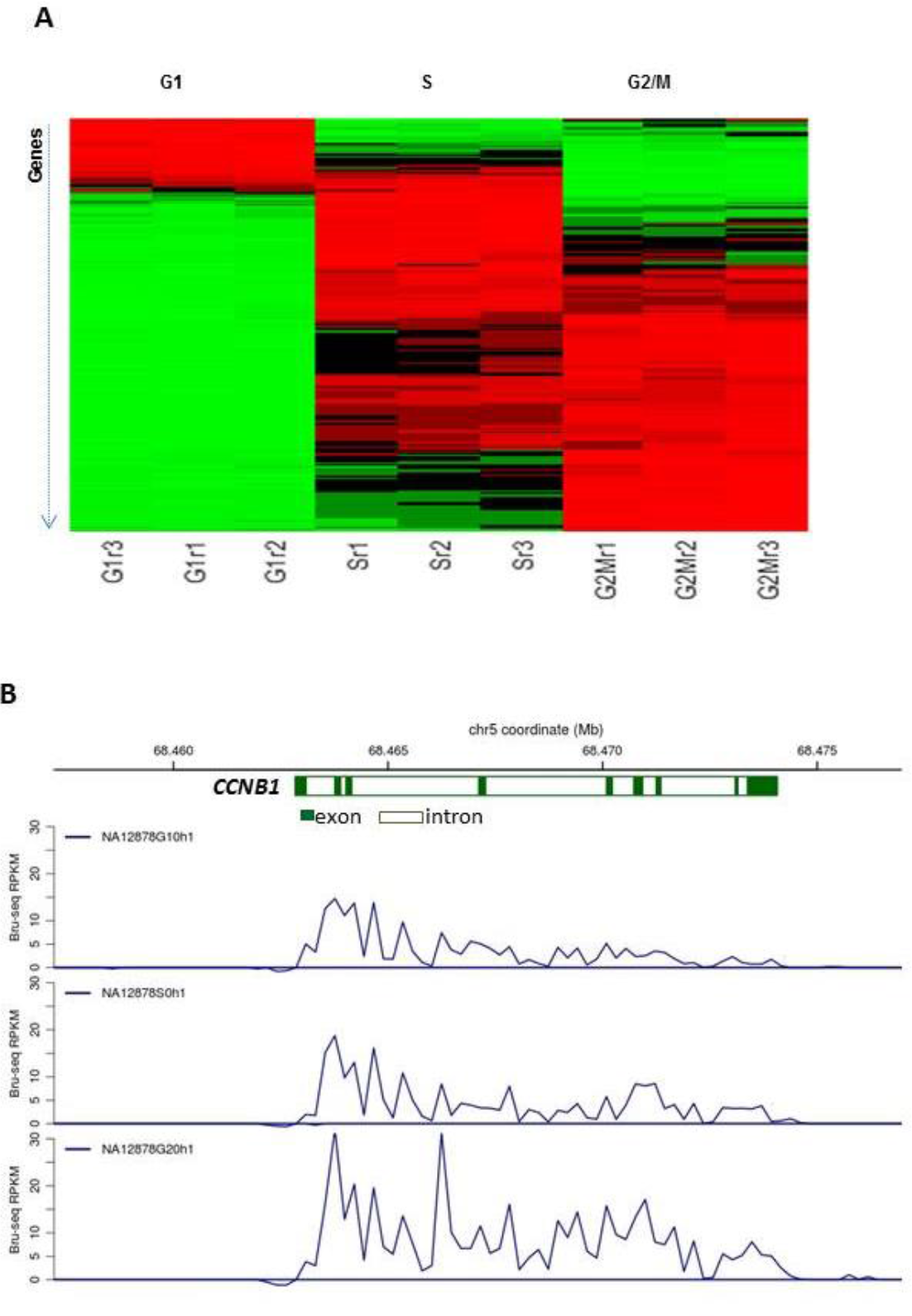
Overview of gene expression for NA12878 cell populations at cell cycle phases G1, S, and G2/M. A. Bulk data analysis RNA-seq data shows a set of cell cycle regulated genes changing expression levels in different cell cycle stages (RNA-seq data, red color indicates increased expression, and green shows reduced expression at a given cell cycle phase compared to other phases). B. Bru-seq measurement of the nascent transcript of *CCNB1* at G1, S, and G2/M. The Y-axis indicates RPKM values, and the X-axis shows mapped Bru-seq reads along the gene body coordinates.

**Extended Data Figure 2.**
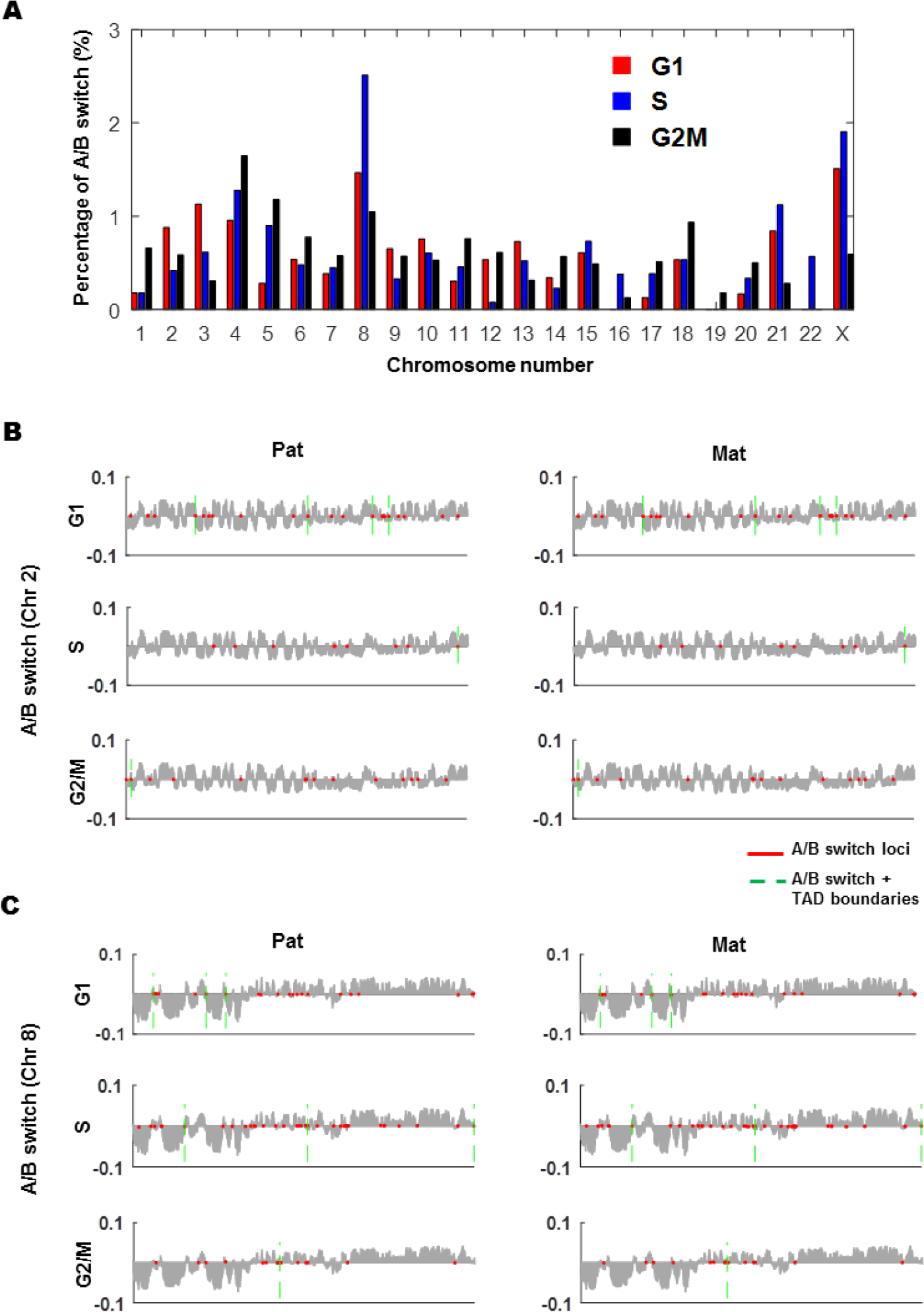
A/B switching regions over the entire genome. (A) Percentage of bins that show changes in A/B compartment status between alleles versus chromosome number for all cell cycle phases, only 0.2 – 2.7% of the genome having A/B switch. (B) A/B compartment pattern of chromosome 3 for each cell cycle stage. (C) A/B compartment pattern of chromosome 8 for each cell cycle stage.

**Extended Data Figure 3.**
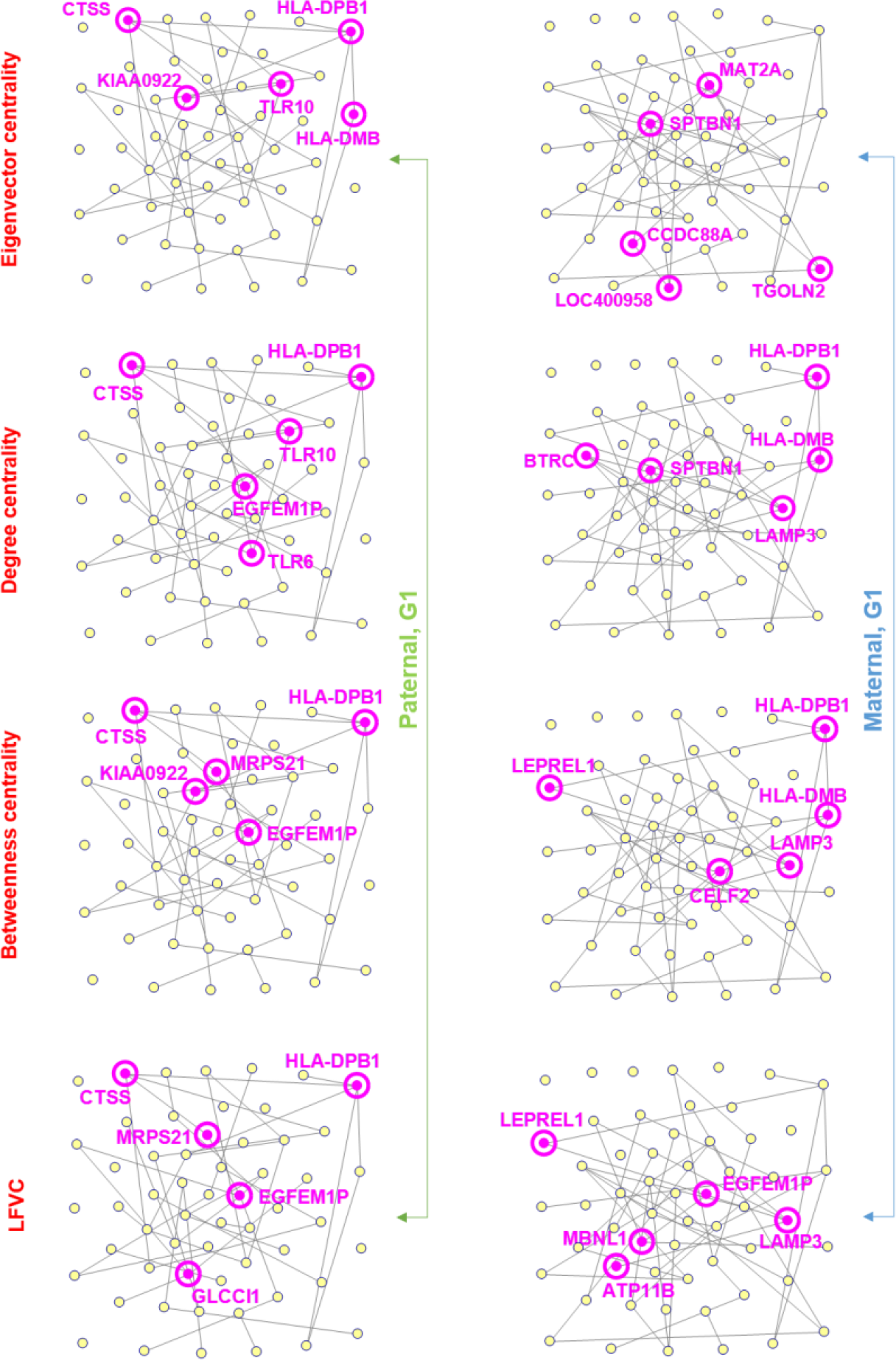
Illustration of network centrality features. Example network is built on the inter-gene contact map at G1 for 57 allelically biased genes within ch1~8. The considered centrality measures include eigenvector centrality, degree centrality, betweenness centrality, and local Fiedler vector centrality (LFVC). Genes with the top 5 largest centrality values are highlighted in magenta. Different centrality methods evaluate the nodal importance in the network from different perspectives.

**Extended Data Figure 4.**
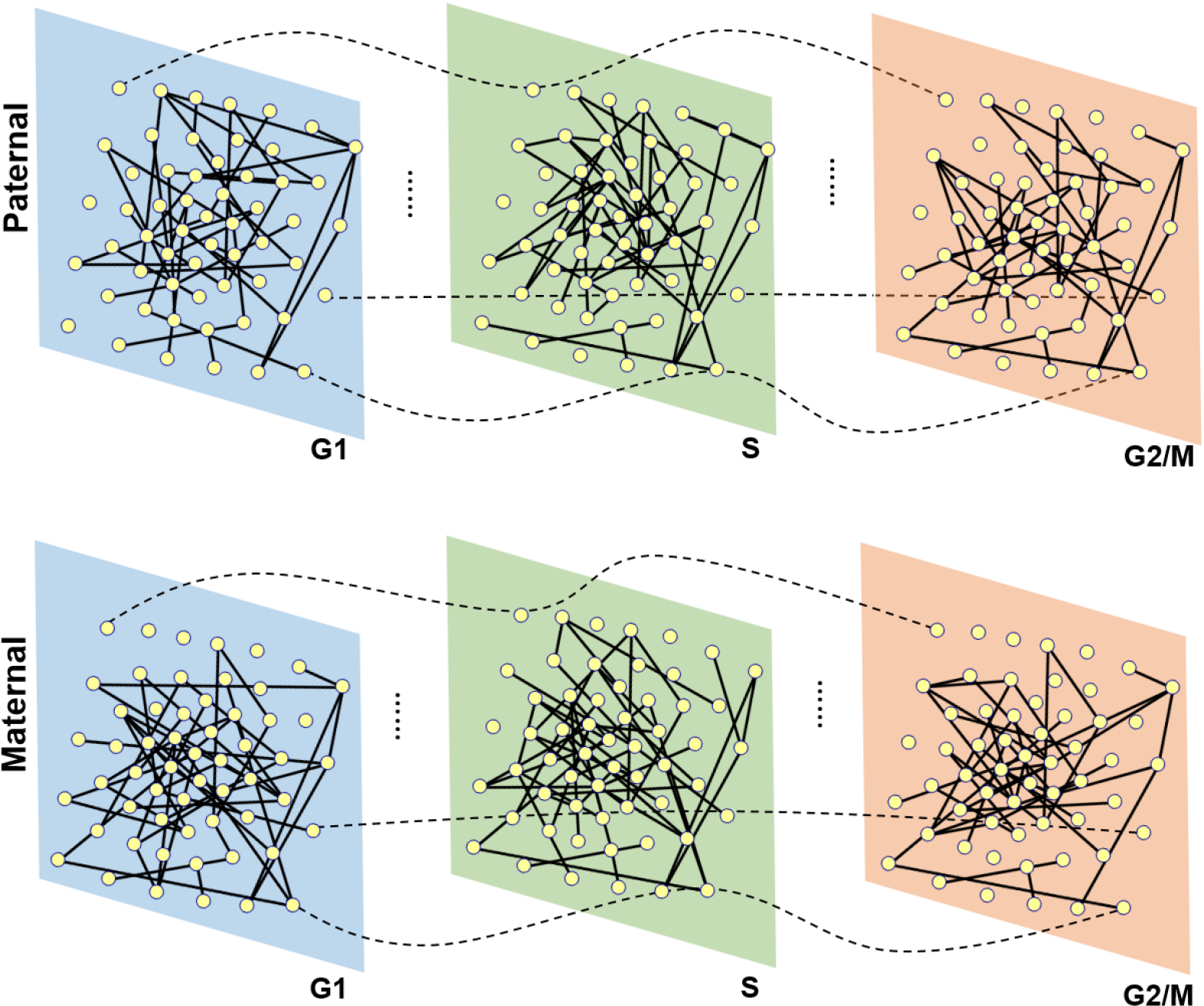
Schematic illustration of multilayer network representation of inter-gene contact map. Each node represents a gene shown in Extended Data Figure 4, and each layer represents a cell cycle phase. The inter-layer connection (dash edge) associates one gene to its counterpart in layers before and after the present layer. The multilayer network representation models the temporal network as an unification.

**Extended Data Figure 5.**
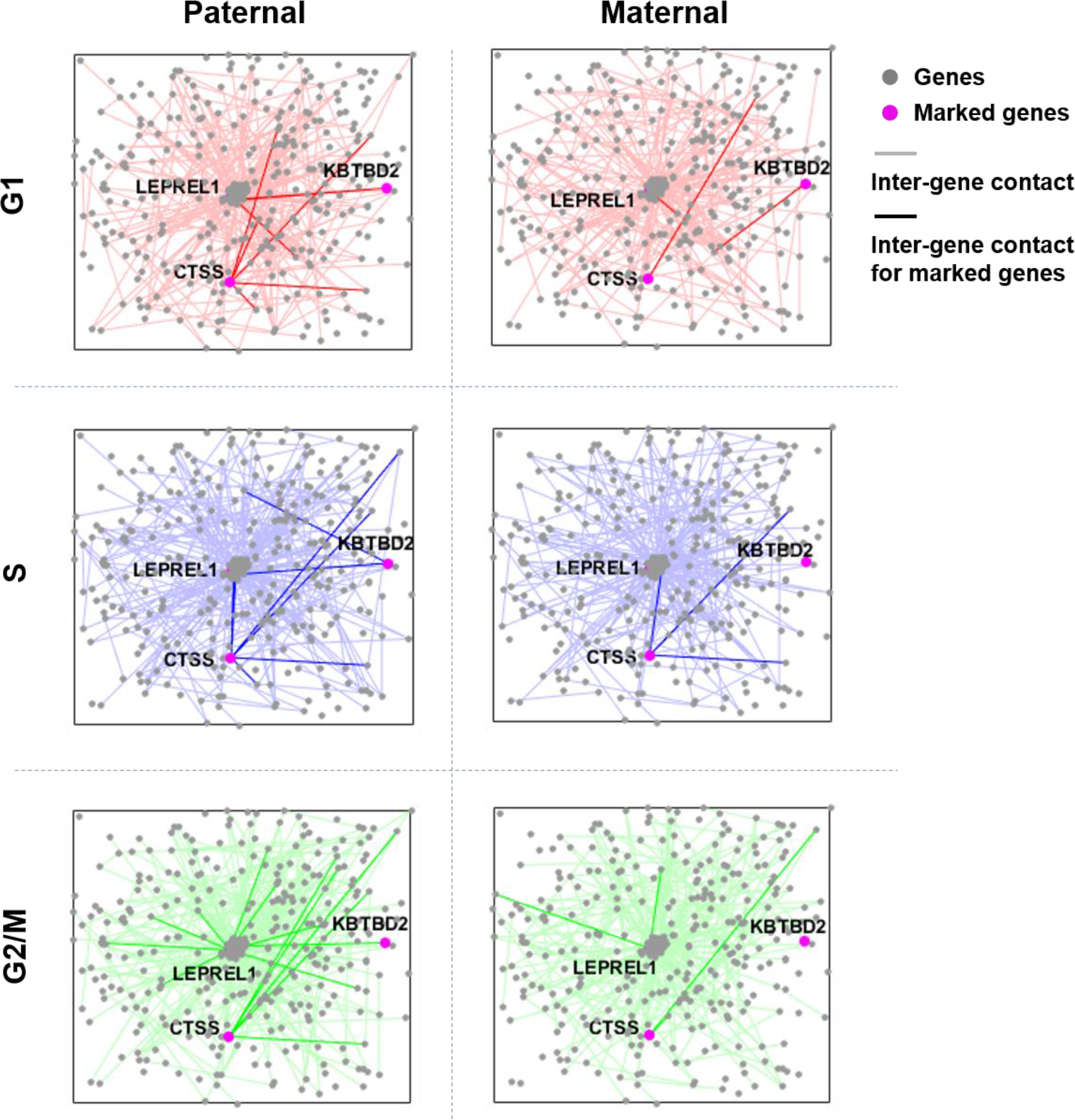
Allele-specific inter-gene contact networks during the cell cycle.

**Extended Data Figure 6.**
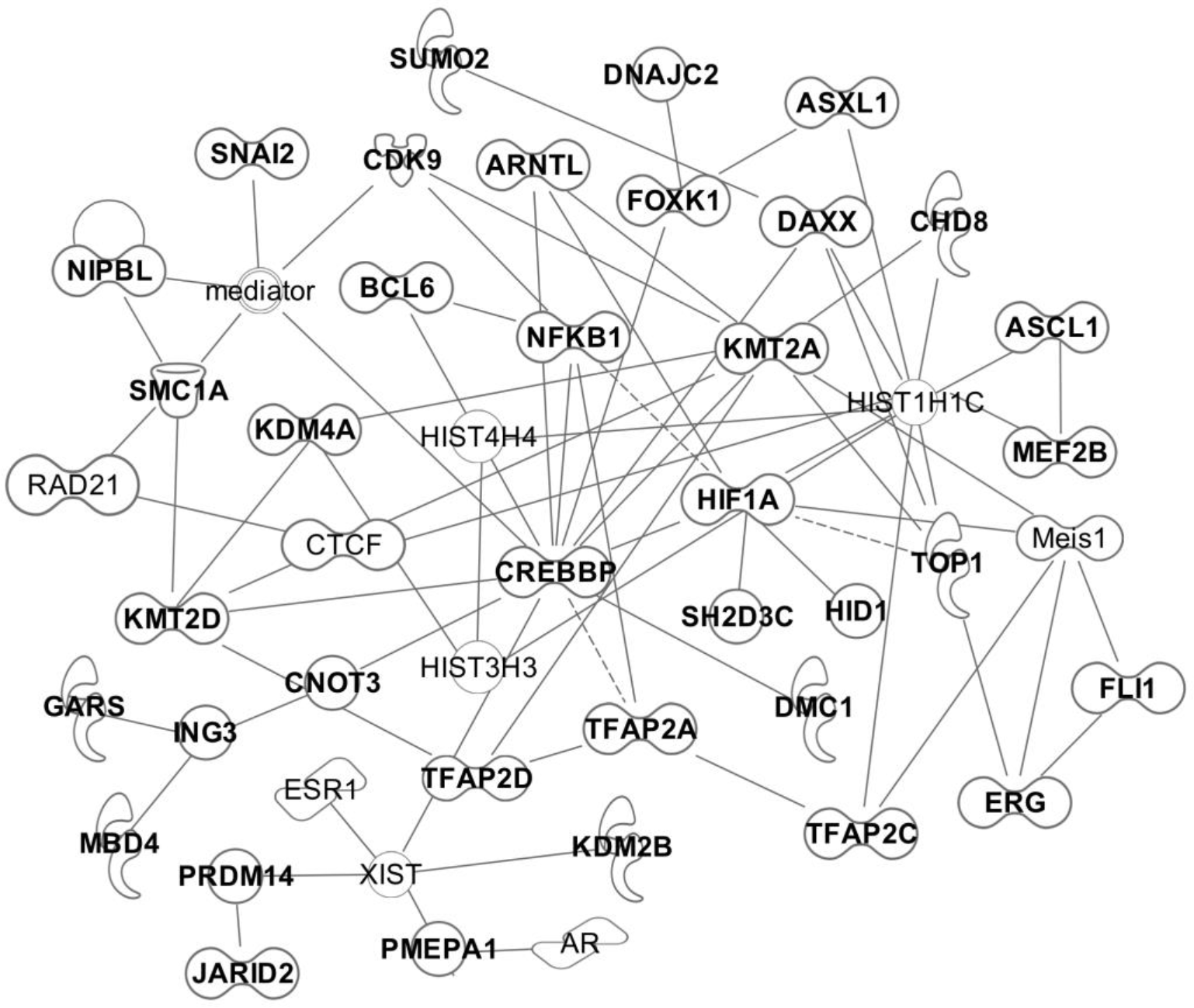
Pathway containing the consensus transcription factor subset (bolded font) bound to the ABE genes. Pathway generated using IPA^®^ ^32^ using filters and the grow function. Additional members of this pathway are not in bold font, and include AR (androgen receptor), CTCF (CCCTC-binding factor), ESR1 (estrogen receptor 1), HIST1H1C (histone cluster 1 H1 family member c), HIST3H3 (histone cluster 3 H3), HIST4H4 (histone cluster 4 H4), the mediator complex, Meis1 (Meis homeobox 1), RAD21 (RAD21 cohesin complex component) and XIST (X inactive specific transcript (non-protein coding).

## Extended Data Table Titles

Extended Data Table 1, List of significant genes from Bulk RNA-seq pair-wise analysis

Extended Data Table 2, List of significant genes from Bulk Bru-seq pair-wise analysis

Extended Data Table 3, Common significant genes Bulk Bru-seq and RNA-seq pair-wise analysis

Extended Data Table 4, Functional annotation of significant genes identified from RNA-seq analysis

Extended Data Table 5, List of ABE genes from RNA-seq analysis

Extended Data Table 6, List of ABE genes from Bru-seq analysis

Extended Data Table 7, Mono-allelic genes identified from RNA-seq analysis

Extended Data Table 8, Genome locations of A/B compartment switching

Extended Data Table 9, Top 10% TADs with allelic difference between the Pat and Mat genomes

Extended Data Table 10, Top 10% genes with inter-gene Hi-C contact difference between the Pat and Mat genomes

Extended Data Table 11, List of genes mapped near TAD boundaries

Extended Data Table 12, Transcription factor with binding site enriched in ABE genes mapped to TAD boundaries.

